# Gene Age Gap Estimate (GAGE) for major depressive disorder: a penalized biological age model using gene expression

**DOI:** 10.1101/2024.09.03.610913

**Authors:** Yijie (Jamie) Li, Rayus Kuplicki, Bart N. Ford, Elizabeth Kresock, Leandra Figueroa-Hall, Jonathan Savitz, Brett A. McKinney

## Abstract

Recent associations between Major Depressive Disorder (MDD) and measures of premature aging suggest accelerated biological aging as a potential biomarker for MDD susceptibility or MDD as a risk factor for age-related diseases. Residuals or “gaps” between the predicted biological age and chronological age have been used for statistical inference, such as testing whether an increased age gap is associated with a given disease state. Recently, a gene expression-based model of biological age showed a higher age gap for individuals with MDD compared to healthy controls (HC). In the current study, we propose an approach that simplifies gene selection using a least absolute shrinkage and selection operator (LASSO) penalty to construct an expression-based Gene Age Gap Estimate (GAGE) model. We train a LASSO gene age model on an RNA-Seq study of 78 unmedicated individuals with MDD and 79 HC, resulting in a model with 21 genes. The L-GAGE shows higher biological aging in MDD participants than HC, but the elevation is not statistically significant. However, when we dichotomize chronological age, the interaction between MDD status and age has a significant association with L-GAGE. This effect remains statistically significant even after adjusting for chronological age and sex. Using the 21 age genes, we find a statistically significant elevated biological age in MDD in an independent microarray gene expression dataset. We find functional enrichment of infectious disease and SARS-COV pathways using a broader feature selection of age related genes.

## 1. Introduction

Major depressive disorder (MDD) has been hypothesized to show characteristics of premature aging (Ford and Savitz (2022)). Biological aging can be measured in multiple dimensions such as telomere length, immunosenescence, brain volume, and gene expression. These measures of biological aging are correlated with chronological age, but environmental and genetic factors can increase or decrease an individual’s biological age relative to their chronological age and influence their risk for age related diseases. For example, MDD has been associated with markers of cellular and immune aging including shortened leukocyte telomere length (Darrow et al., 2016; Ridout et al., 2016), elevated indicators of oxidative stress (Ait Tayeb et al., 2023), and elevated circulating inflammatory cytokines (Raison et al., 2006). Epigenetic clocks predicting biological age based on the accumulation of methylated CpG sites have found higher biological age in MDD participants compared with healthy controls (Protsenko et al., 2021). Brain age models constructed from T1-weighted magnetic resonance image (MRI) data from 2,188 healthy controls predicted a gap of +1.08 years (SE 0.22) between predicted and chronological age across 2,675 depressed participants (Han et al., 2021).

A recent RNA-Seq MDD study found that gene expression-based biological aging was elevated in MDD participants compared to Healthy Controls (HC) (Cole et al., 2021). The PBMC samples included four groups: 44 healthy controls and a mixture of MDD participants: 94 treatment-resistant, 47 treatment-responsive, and 46 untreated (Cole et al., 2021). They selected age genes iteratively by varying the P-value threshold for the t-test between upper and lower chronological age quartiles. For a given iteration, a biological age was computed for each subject based on the signed z-score of the age-related genes, and the P-value threshold was chosen to optimize the correlation between biological and chronological age of the participants (Spearman Correlation Coefficient (SCC) = 0.72, p < 0.01). A linear model of biological age was fit to chronological age and association with MDD was computed by comparing the number of MDD and HC participants above and below the regression line.

In the current study, we describe a different gene age approach that simplifies the gene selection procedure by modeling age from RNA-Seq gene expression using a multivariate LASSO penalized regression rather than an iterative univariate test. The LASSO approach has the potential to capture more variation because it is multivariate, plus it automates gene selection by cross-validation of the penalty and it reduces the amount of correlation in the selected features. We use age as a quantitative variable during the LASSO feature selection in linear regression, as opposed to using age quartiles, which is another way for the model to include more variation when estimating age. After training the gene age model, we dichotomize chronological age when using it as a covariate for the association of the gene age gap with MDD.

The current study is outlined as follows. We describe the LASSO biological age model trained on an existing RNAseq dataset consisting of 157 individuals (78 with MDD and 79 healthy controls) (Li et al., 2022). The residual is an estimate of the gap between an individual’s chronological age and their biological gene age, which we refer to as the LASSO Gene Age Gap Estimate (L-GAGE). We describe the use of L-GAGE for testing elevated biological aging in MDD. We find L-GAGE is elevated in MDD participants compared to HC, but the elevation is not statistically significant. However, when we dichotomize chronological age into older and younger, the interaction between MDD status and age is significantly associated with the L-GAGE residual. We use the top L-GAGE genes to train a gene age model in an independent public dataset for MDD, and the residual shows a statistically significant increase in MDD compared to HC. Finally, we use machine learning feature selection to explore biological pathways that are significantly enriched for the gene sets identified as being associated with aging.

## 2. Materials and Methods

### 2.1. Gene Expression Data

To train our biological age models, we use an extant RNA-Seq dataset (Le et al., 2020). The study was approved by the Western Institutional Review Board and conducted according to the principles expressed in the Declaration of Helsinki. The data consists of 78 MDD and 79 HC participants (91 females and 66 males). Individuals with current symptoms of depression met DSM-IV-TR criteria for MDD based on the Structural Clinical Interview for DSM-IV-TR Axis I Disorders and an unstructured psychiatric interview. HC individuals had no personal or immediate family history of major psychiatric disorders. MDD participants were unmedicated for at least 3 weeks prior to study entry. Exclusion criteria included major medical or neurological illness, psychosis, traumatic brain injury, and a history of drug/alcohol abuse within 1 year. There is a higher female/male ratio for MDD (51/27) than HC (40/39), compatible with trends in the general population. The age distribution is slightly skewed towards younger individuals with age range from 18 to 55 (Fig. 1). The 8,923 genes in the RNA-Seq gene expression data are normalized by counts per million reads, which we then quantile normalize and log2 transform to stabilize variance. We removed genes with a low coefficient of variation (standard deviation divided by absolute mean). We chose a threshold of 0.045 to obtain 5,587 genes.

**Figure 1.**
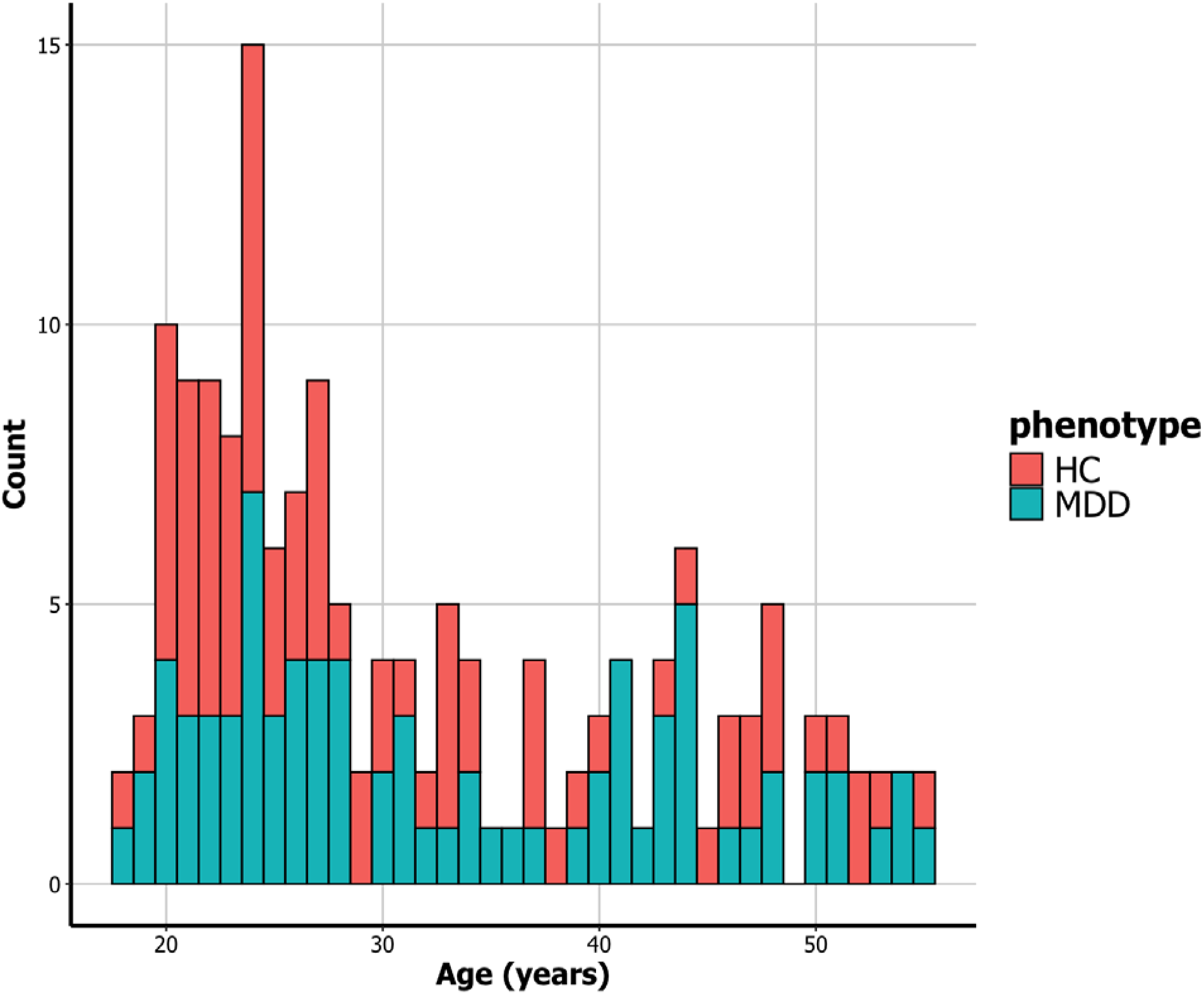
Histogram of chronological ages with a bin size of 1: Bars are separated by Healthy Control (HC, red) and major depressive disorder (MDD, blue). There are more younger participants in the dataset with the same age, especially from age 20∼28. For example, there are 15 participants that are 24 years of age. Chronological age is not associated with MDD versus HC (T-test P-value 0.167).

To test the generalizability of the gene age model, we use an independent microarray study of MDD from the gene expression omnibus (GEO) with accession number GSE98793 (Leday et al., 2018). This data skews older than the discovery data (ages ∼30–70 years) and includes MDD with anxiety. We exclude anxiety, resulting in 64 participants with MDD and 64 HC.

#### Gene Age Gap Estimate (GAGE)

We use LASSO for gene selection and modeling biological age, and then we use the residual of this model, which we call LASSO Gene-Age Gap Estimate (L-GAGE), for association testing with MDD. For the LASSO biological aging model, we build a full penalized regression model with all gene expression variables and with chronological age as the outcome variable. We include both MDD and HC samples in the age model, which was also the approach in Ref. (Cole et al., 2021). Our biological age model is based on the non-zero coefficient genes from the lambda-1se LASSO penalty (the largest A for which the average cross-validation (CV) error is within one standard error of the minimum CV error). We compute the gap/residuals of the LASSO model between predicted biological age and chronological age (i.e., the L-GAGE score). Our goal is to use L-GAGE to test for increased biological age in MDD participants (Fig. 2).

**Figure 2.**
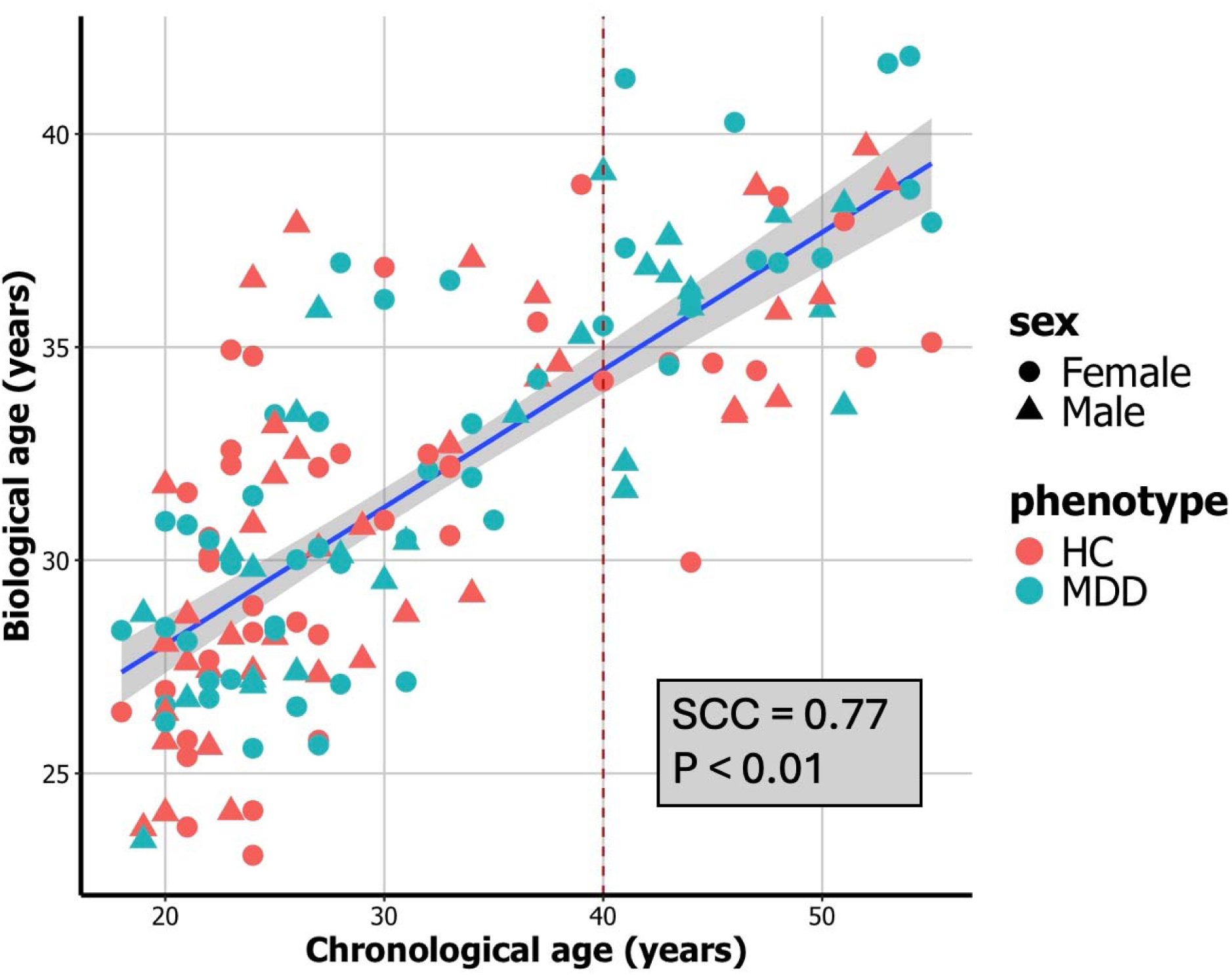
Scatter plot with regression line of biological age and chronological age: Biological age model is based on LASSO regression and the residual is later used for LASSO Gene Age Gap Estimate (L-GAGE). The points are colored by MDD (blue) and HC (red). The points are shaped by Female (circle) and Male (triangle). Spearman Correlation Coefficient (SCC = 0.77, slope P-value < 0.01).

For the replication microarray dataset, we retrain the gene-age regression coefficients for the top L-GAGE genes because we could not match two of the gene symbols that are part of the multivariate L-GAGE model from the RNA-Seq data. We use Ridge rather than LASSO because we first perform feature selection in the discovery data and LASSO may force too many genes to zero. We test Ridge residual (R-GAGE) for association with MDD in the microarray data.

### 2.2. Relationship between gene age gap, chronological age, MDD and sex

It is important to consider adjustments for chronological age in biological age models because of regression to the mean as discussed for brain age models (Le et al., 2018), but sex is also an important covariate for MDD. To further explore covariate effects, we add MDD x Age and MDD x Sex interactions for L-GAGE associations with MDD. We use the OLS model

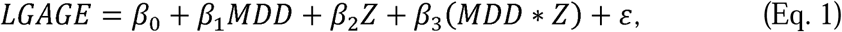

where Z represents the adjustment or interaction variable (Age or Sex). We focus on the effect of β_3_, which represents how much the average L-GAGE of the MDD group changes for the Z=1 condition.

We consider two cases when age is used as a covariate with interactions (Z in Eq. 1): as continuous and as dichotomous with a threshold. To verify our choice of age threshold, we use a threshold regression model in the “chngpt” package in R (Fong et al., 2017). We use this approach to check for possible nonlinear relationship between MDD and age and whether the effect of chronological age on MDD increases at some threshold point. The mean function of the threshold model is:

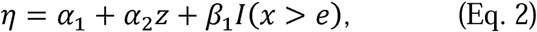

where x stands for chronological age, e is the age threshold and z are additional predictors. “I” is a step indicator function. The threshold is optimized using the exact criterion function with a logistic-based smooth function.

### 2.3. Feature selection, Gene-Age pathway Enrichment, and interpretable classifier

We use LASSO to create the gene-based residual age model, L-GAGE, but LASSO feature selection also results in a set of age-related genes. As a secondary analysis, we use LASSO and other feature selection methods to identify important age-related genes for pathway enrichment to understand the biological mechanisms of the age models. We use univariate linear regression, random forest (RF) regression, and nearest-neighbor projected distance regression (NPDR) (Le et al., 2020) as feature selection methods. RF has the ability to find more complex models than LASSO and linear regression, but RF has limited ability to detect interactions (McKinney et al., 2009), whereas NPDR has the ability to detect interaction effects (Le et al., 2020). For univariate feature selection, we use a linear model of individual genes with age, and we use a P-value threshold of 0.05 (uncorrected for improved pathway overlap). We use the standard NPDR with an adjusted P-value threshold of 0.05 FDR, and we use the LASSO penalized NPDR. For NPDR, we use the imbalanced k-nearest-neighbor value (k=47) that approximates the 0.5 standard deviation of the hyper-radius (Le et al., 2020). We use permutation variable importance with RF. We use the Reactome Pathway database in MSigDB (Subramanian et al., 2005) for biological pathway enrichment of age related genes.

For additional interpretation of the gene-age prediction of MDD along with consideration for other covariates, we train a decision tree to predict MDD based on L-GAGE, chronological age, and sex. Decision trees have high variance, but they are useful for interpreting the relationships between covariates.

## 3. Results and Discussion

### 3.1. Association Testing of Gene Age L-GAGE with MDD

We test for association of the LASSO Gene Age Gap Estimate (L-GAGE) score with MDD status. L-GAGE is the residual from a LASSO gene expression model of chronological age. The LASSO model uses the cross-validation tuned lambda-1se value (A *= 1.636048*), which is the largest A at which the mean-squared error (MSE) is within one standard error of the minimum MSE. The residuals are constant, and heteroscedasticity is not present based on the Non-constant Variance Score Test. The penalty results in a multivariate linear model of age with 22 genes and a Spearman Correlation Coefficient (SCC) with chronological age of 0.77 (Fig. 2). Counting the number of HC or MDD above or below the regression line (Fig. 2), we find that the biological age is greater in MDD participants than HC (HC – 45 (56.96 %) below, 34 (43.037%) above, MDD 35 (44.87%) below, 43 (55.128%) above). The P-value of the Chi-squared test of GAGE sign (above or below the line) for MDD is not significant (0.1753). The greater L-GAGE in MDD versus HC can be seen in L-GAGE density (Fig. 3A). The L-GAGE distribution for males and females is very similar (Fig. 3B). While L-GAGE is greater in MDD than HC participants, we do not find a statistically significant replication of the effect found in Ref. (Cole et al., 2021). However, we do see a comparable effect size to what was previously found. Using the same genes as their model also did not yield a statistically significant MDD association.

**Figure 3.**
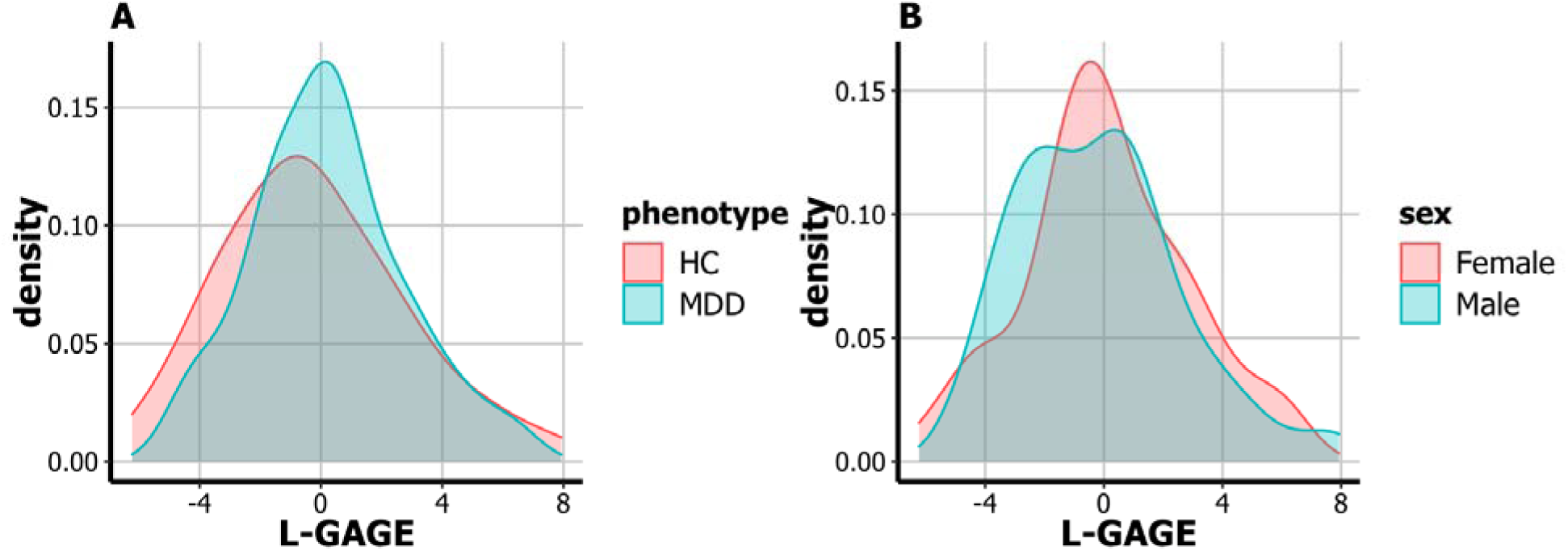
Density plots of the LASSO based Gene Age Gap Estimate (L-GAGE) separated by MDD (A) and sex. **(B)**. A positive gene-age residual (x-axis) indicates a sample above the gene age regression line and negative below. **A**. Biological age relative to chronological age (L-GAGE) is greater in MDD participants than in HC. **B**. The L-GAGE difference between males and females is less pronounced.

### 3.2 Testing MDD-Age interaction for L-GAGE association model

We test for the effect of L-GAGE on MDD by introducing an MDD-Age interaction term (Eq. 1). Dichotomizing age at threshold 40, MDD alone is not significant, but we find a statistically significant effect of the interaction between MDD and Age 40 on L-GAGE (Table 1 and Fig. 4). For individuals younger than 40, L-GAGE shows very little difference between MDD and HC, but for older individuals, there is greater biological aging (L-GAGE) for the MDD versus HC group (Table 1 and Fig. 4). Age alone is also statistically significant (Table 1). These age effects remain significant when we add sex as a covariate (Table 1B), but sex is not significant (Table 1B and Table 2).

**Figure 4.**
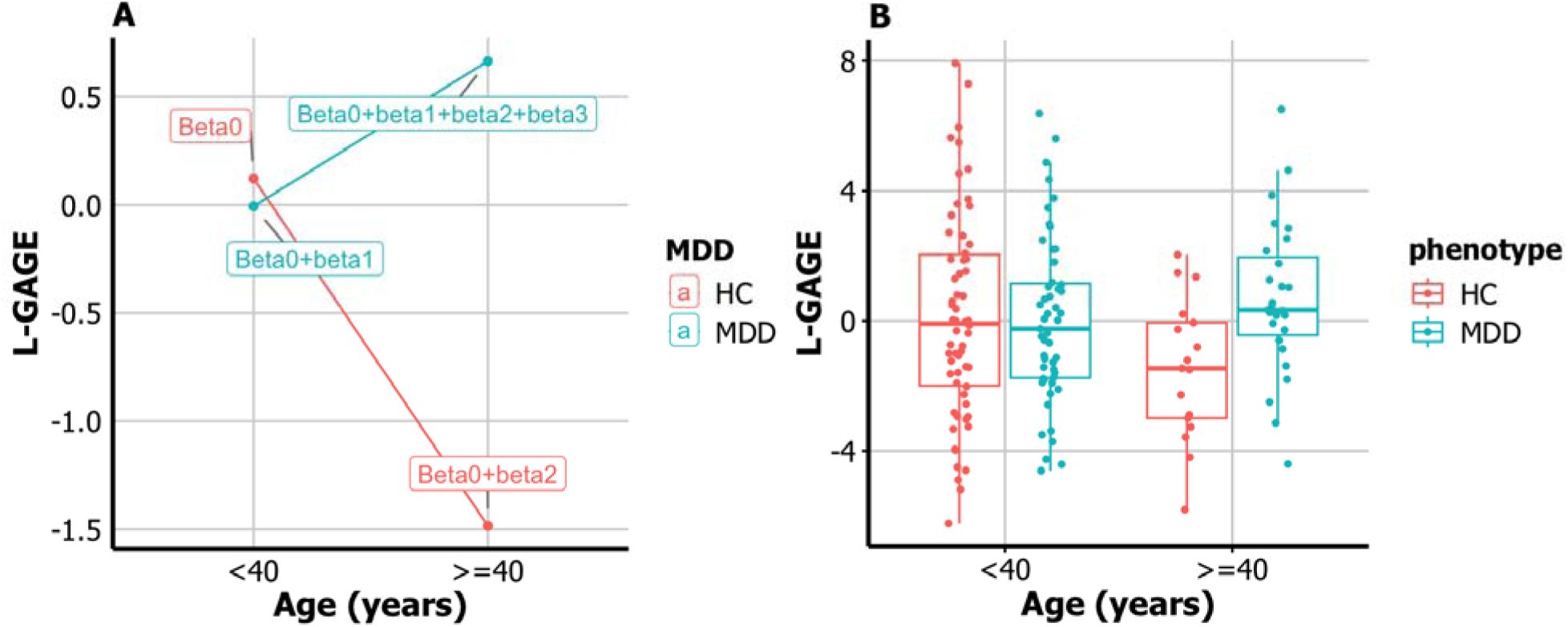
MDD x Age interaction for L-GAGE with age 40 threshold. **A.** The average L-GAGE for people older than 40 with MDD is higher than the L-GAGE value for people younger than 40 with MDD (blue line), whereas in the HC group the average L-GAGE is lower for people older than 40 than for people younger than 40 (red line). **B.** For individuals younger than 40, L-GAGE shows very little difference between MDD and HC. For older individuals, there is greater biological aging (L-GAGE) for the MDD versus HC group. The L-GAGE association with MDD is still significant when adjusted by age and sex.

**Table 1.** LASSO Gene-age-gap estimate (L-GAGE) association with MDD and dichotomized age interaction.

|  | Estimate | Std. Error | t value | Pr(>\|t\|) |
| --- | --- | --- | --- | --- |
| (Intercept) | 0.1225 | 0.3495 | 0.35 | 0.7265 |
| MDD | -0.1286 | 0.5203 | -0.247 | 0.8052 |
| Age40 | -1.606 | 0.7535 | -2.131 | 0.0347* |
| MDD*Age40 | 2.2764 | 0.9984 | 2.28 | 0.024* |
| Signif. codes: 0 '***' 0.001 '**' 0.01 '*' 0.05 '.' 0.1 ' ' 1 |  |  |  |  |

B
|  | Estimate | Std. Error | t value | Pr(>\|t\|) |
| --- | --- | --- | --- | --- |
| (Intercept) | 0.2644 | 0.4172 | 0.634 | 0.5273 |
| MDD | -0.1870 | 0.5296 | -0.353 | 0.7246 |
| SexMale | -0.2838 | 0.4534 | -0.626 | 0.5324 |
| Age40 | -1.6144 | 0.7551 | -2.138 | 0.0341* |
| MDD*Age40 | 2.3274 | 1.0038 | 2.319 | 0.0217* |
| Signif. codes: 0 '***' 0.001 '**' 0.01 '*' 0.05 '.' 0.1 ' ' 1 |  |  |  |  |

*Note. A.* Based on the ordinary least squares model (Eq. 1 with Z=Age), where chronological age is dichotomized with threshold is Age ≥ 40 and Age $<$ 40, the MDD x Age interaction is significant. Biological age (L-GAGE)) is similar for MDD and HC when Age $<$ 40, but when the chronological age is higher than 40, biological age is significantly greater in MDD individuals than HC. *B.* The MDD x Age interaction remains significant when Sex is added as a covariate.

**Table 2.** Gene-age-gap regression with MDD-sex interaction with Female and Male.

|  | Estimate | Std. Error | t value | Pr(>\|t\|) |
| --- | --- | --- | --- | --- |
| (Intercept) | -0.24481 | 0.44221 | -0.554 | 0.581 |
| MDD | 0.64637 | 0.5907 | 1.094 | 0.276 |
| Male | 0.04395 | 0.62938 | 0.07 | 0.944 |
| MDD* Male | -0.55121 | 0.91608 | -0.602 | 0.548 |
| Signif. codes: 0 '***' 0.001 '**' 0.01 '*' 0.05 '.' 0.1 ' ' 1 |  |  |  |  |

*Note.* Based on the ordinary least squares model (Eq. 1 with Z=Male/Female instead of age), L-GAGE score of MDD in males is slightly lower than the L-GAGE score of MDD in females, but the interaction term MDD\*Male is not statistically significant.

The MDD-Age interaction and the MDD term (Eq. 1) do not have a significant effect on L-GAGE when age is treated as a continuous variable (MDD P-value = 0.364, Age P-value = 0.316, MDD*Age P-value = 0.197). Also, there is no direct statistical association between MDD and age and between MDD and sex (Two Sample T-test of MDD and Chronological age: P-value = 0.167; Chi-squared-test of MDD and sex: P-value = 0.08716). To further support our choice of age threshold, we use a threshold regression (Eq. 2). The change point for age in relation to MDD is estimated to be 39 years (Fig. 5). Combined with the third quartile being age 41, the threshold regression suggests that age 40 is a suitable cutoff point for dividing the participants into two age groups.

**Figure 5.**
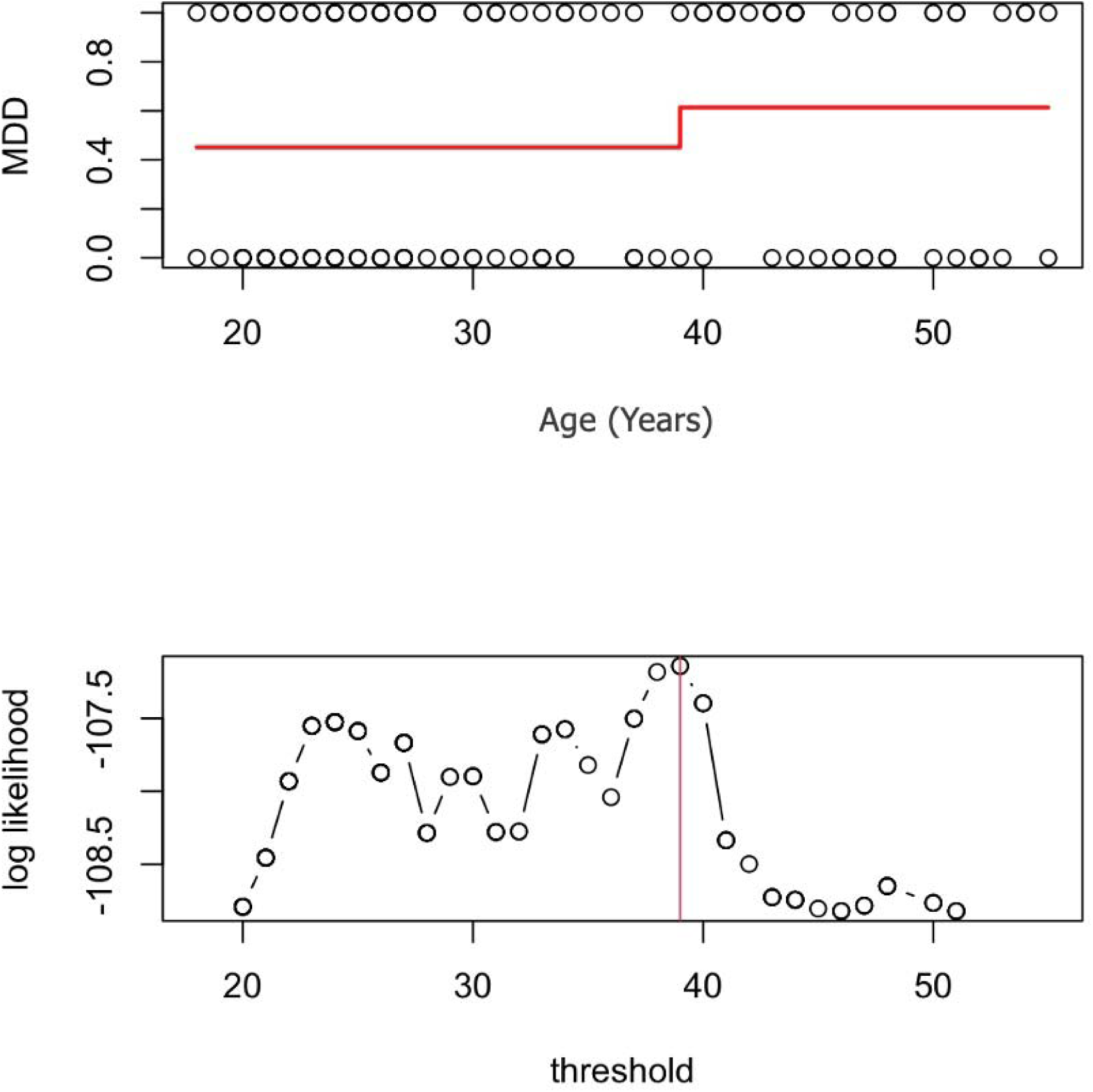
Effect of Chronological Age on MDD Determined by Threshold Regression Model. **A.** Threshold regression (Eq. 2) shows the nonlinear relationship between MDD and chronological age. The prediction indicates an increase in MDD up to the age of 39, which is identified as the change point by the model. **B.** The likelihood analysis of the threshold regression model also indicates that age 39 is the optimal threshold, having the highest model likelihood.

Additional support for the age-40 threshold can be seen in the decision tree for predicting MDD (Fig. 6), where age with threshold 39.5 is the second important split variable, following L-GAGE. The decision tree also suggests interaction effects, where the effect of L-GAGE on MDD is conditioned on chronological age. If L-GAGE (node 1, Fig. 6) is below a threshold, participants tend to be HC. If the L-GAGE is below the threshold and chronological age is above 39.5 (i.e., an interaction), participants tend to be MDD. However, for chronological age less than 39.5. (node 3, Fig. 6), the prediction of MDD is considerably more complex. We note that this decision tree was trained on the full dataset to maximize power, but it is instructional for interpretation.

**Figure 6.**
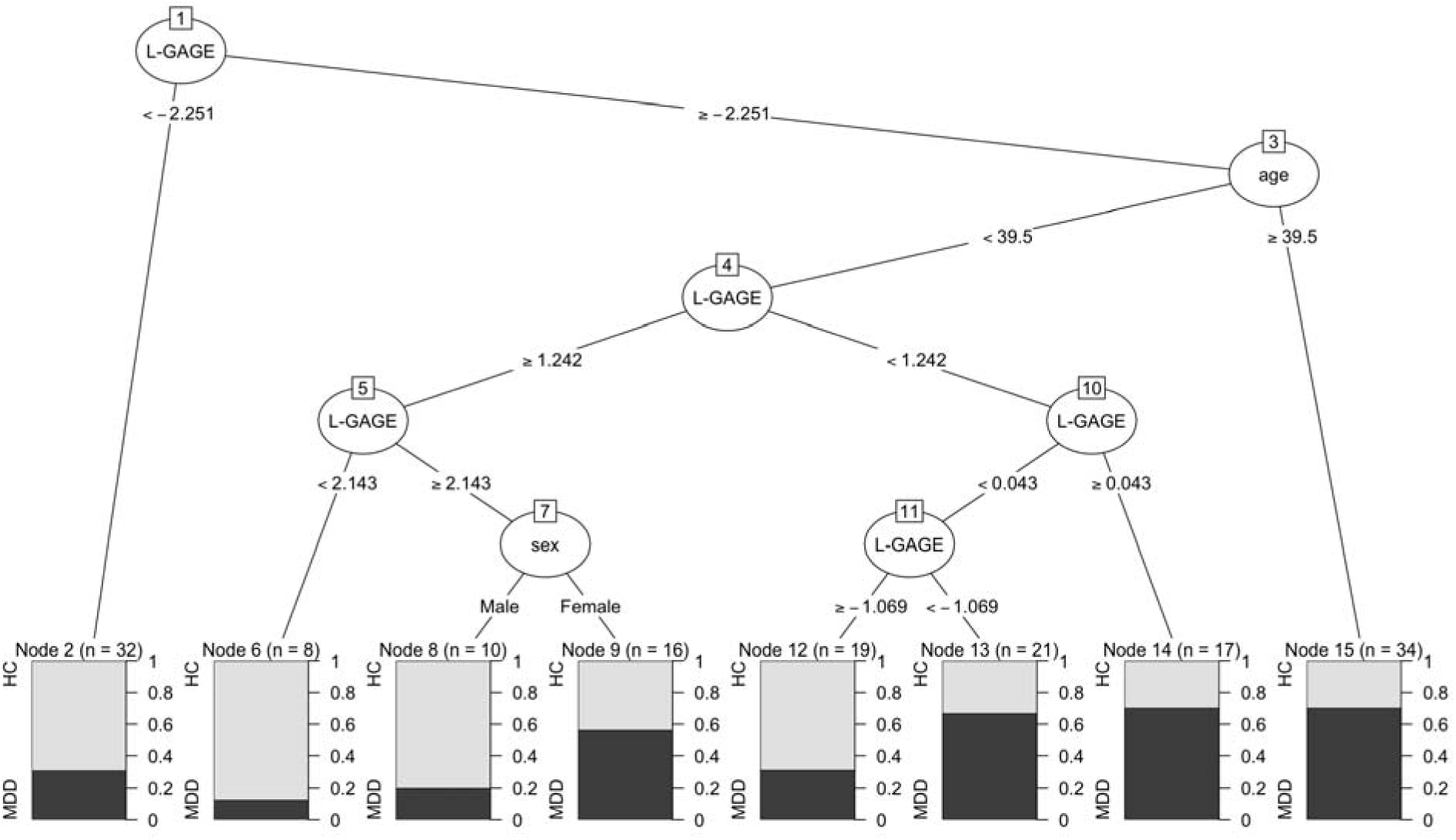
Gene age decision tree for MDD with covariates. For added interpretation, we train a decision tree on all samples to predict MDD. The model identifies the gene age residual L-GAGE as the most important predictor, with chronological age being the second most significant factor. In the first split, if the gene age gap is low, L-GAGE < −2.251 (Node 1), there is high probability for a subject to be HC (Node 2). If the gene age gap is higher, L-GAGE ≥ −2.251, the model becomes more complex and initially depends on chronological age with split 39.5 years (Node 3). If L-GAGE is high and Age ≥ 39.5, then there is a high probability a subject is MDD (Node 15). When Age < 39.5, the model again becomes dependent on L-GAGE, and at a certain split, females exhibit a higher probability of MDD compared to males (Nodes 8 and 9).

### 3.3 Characterizing Age-Associated Genes

The LASSO regression used in L-GAGE selected 21 age genes with non-zero coefficients (Table 3). In order to perform pathway enrichment for age related genes, we expand the number of genes and feature selection methods to include linear regression, RF, and nearest-neighbor projected distance regression (NDPR) (Le et al., 2020). Using the Reactome database, we find enrichment for Infectious Disease, Adaptive Immune System, and SARS-CoV-2 Infection pathways for top genes with P-value<0.05 from linear regression (Table 5) and NPDR (Table 6). SARS-CoV-2 can cause neurological complications, and a recent study showed that differentially expressed genes for COVID infection overlap with many gene associations for neuropsychiatric disorders including depression (Quincozes-Santos et al., 2021). We also found across all feature selection methods (including LASSO), the four common age genes are NAA20 (N-alpha-acetyltransferase 20), CCNE1 (Cyclin E1), and SESTD1 (SET domain containing protein 1A), and TAF9 (TATA-box-binding protein associated factor 9). These genes will be discussed further.

**Table 3.** Age associated genes selected by LASSO.

| Down Regulated with Increasing Age |  | Up Regulated with Increasing Age |  |
| --- | --- | --- | --- |
| Gene | Coefficient | Gene | Coefficient |
| NAA20 | -6.7070152 | CCNE1 | 14.2689027 |
| ZNF347 | -2.9514771 | SESTD1 | 8.8624231 |
| PRMT6 | -2.4559818 | ZNF334 | 2.4209761 |
| WDR13 | -1.7979357 | ANTXRL | 2.0255277 |
| DDX19B | -1.3737037 | DTD2 | 1.8502139 |
| TAF9 | -1.2672137 | CYTH3 | 1.5349361 |
| ADSS | -1.1724134 | DYRK1A | 1.2905045 |
| TGFBR3 | -1.0316785 | HTATSF1 | 1.078388 |
| SMYD5 | -0.8454683 | SFXN4 | 0.7870119 |
| CISD1 | -0.6212633 | UBE2F-SCLY | 0.2252943 |
| TGIF2-C20orf24 | -0.5057642 |  |  |

*Note.* Multivariate coefficients are shown that survived LASSO penalty. Negative coefficients (left columns) indicate higher expression of the gene tends to occur with younger age. Positive coefficients (right columns) indicate higher expression of the gene tends to occur in older individuals. These genes are used in the gene age model and the L-GAGE residual.

**Table 4.** Age associated genes selected by linear regression with adjusted P-value 0.05 FDR.

| Down Regulated with Increasing Age |  |  |  | Up Regulated with Increasing Age |  |  |  |
| --- | --- | --- | --- | --- | --- | --- | --- |
| Gene | Coefficient | P-value | Adjusted P-value | Gene | Coefficient | P-value | Adjusted P-value |
| NAA20 | -16.1918 | 8.86E-08 | 0.0005 | CCNE1 | 42.8022 | 5.59E-07 | 0.0013 |
| CIART | -22.7969 | 6.87E-07 | 0.0013 | SESTD1 | 12.2045 | 1.19E-05 | 0.0111 |
| TAF9 | -21.2804 | 2.85E-06 | 0.0040 | ITGB1BP1 | 10.7847 | 2.46E-05 | 0.0197 |
| MLXIPL | -20.0949 | 4.55E-06 | 0.0051 | ANTXRL | 13.8739 | 4.23E-05 | 0.0295 |
| TGFBR3 | -17.7019 | 7.91E-05 | 0.0491 |  |  |  |  |

*Note.* Negative coefficients (left columns) indicate higher expression of the gene tends to occur with younger age. Positive coefficients (right columns) indicate that higher expression of the gene tends to occur in older individuals. These genes are shown for comparison but not used in the gene age model.

**Table 5.**
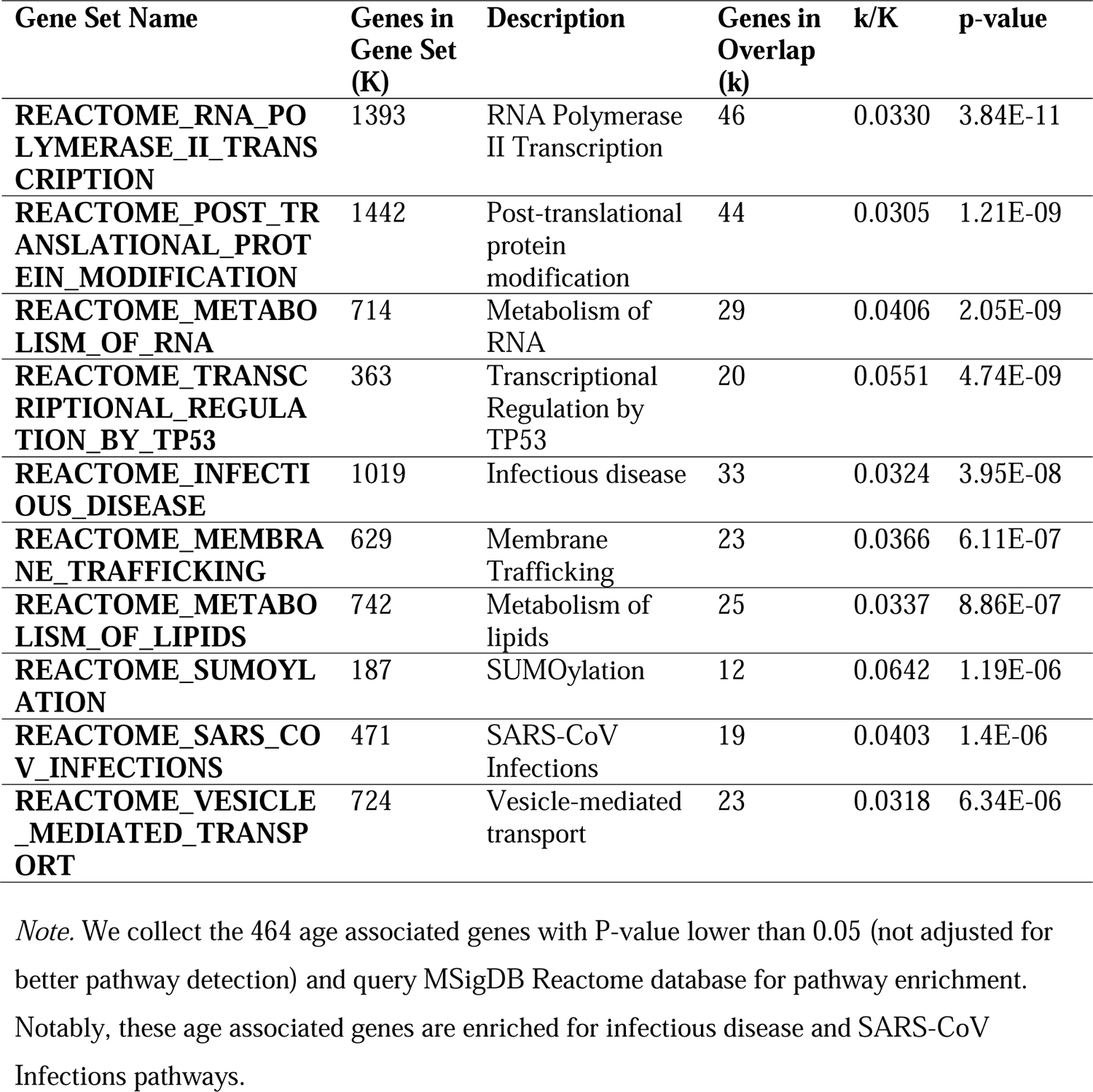
MSigDB Reactome results of the age genes selected by linear regression.

**Table 6.** MSigDB Reactome results of the 145 age genes selected by nearest-neighbor projected distance regression (NPDR) with LASSO penalty.

| Gene Set Name | Genes in Gene Set (K) | Description | Genes in Overlap (k) | k/K | p-value |
| --- | --- | --- | --- | --- | --- |
| REACTOME_NEF_MEDIATES_DOWN_MODULATION_OF_CELL_SURFACE_RECEPTORS_BY_RECRUITING_THEM_TO_CLATHRIN_ADAPTERS | 21 | Nef-mediates down modulation of cell surface receptors by recruiting them to clathrin adapters | 4 | 0.1905 | 7.44E-07 |
| REACTOME_NEF_MEDIATED_CD4_DOWN_REGULATION | 9 | Nef Mediated CD4 Down-regulation | 3 | 0.3333 | 3.22E-06 |
| REACTOME_RNA_POLYMERASE_II_TRANSCRIPTION | 1393 | RNA Polymerase II Transcription | 16 | 0.0115 | 2.38E-05 |
| REACTOME_LDL_CLEARANCE | 19 | LDL clearance | 3 | 0.1579 | 3.62E-05 |
| REACTOME_TRANSCRIPTIONAL_REGULATION_BY_TP53 | 363 | Transcriptional Regulation by TP53 | 8 | 0.022 | 3.78E-05 |
| REACTOME_MHC_CLASS_II_ANTIGEN_PRESENTATION | 126 | MHC class II antigen presentation | 5 | 0.0397 | 7.56E-05 |
| REACTOME_ADAPTIVE_IMMUNE_SYSTEM | 829 | Adaptive Immune System | 11 | 0.0133 | 1.38E-04 |
| REACTOME_TRAFFICKING_OF_AMPA_RECEPTORS | 31 | Trafficking of AMPA receptors | 3 | 0.0968 | 1.63E-04 |
| REACTOME_TP53_REGULATES_METABOLIC_GENES | 87 | TP53 Regulates Metabolic Genes | 4 | 0.046 | 2.32E-04 |

### 3.4 Association Test of L-GAGE in independent MDD study

To evaluate the generalizability of the 21 genes from the L-GAGE model, we train a Ridge regression model of age using a microarray study by (Leday et al., 2018). Out of the 21 genes in our L-GAGE model, 19 were found in the microarray dataset by matching gene symbols (genes ANTXRL and UBE2F-SCLY were not present). Using the 19 genes in the microarray data with 64 MDD and 64 HC, the Ridge model of chronological age (lambda penalty 1.991) has SCC=0.465 (Fig. 7). In the scatter plot of Ridge gene-age versus chronological age (Fig. 7) 41(23) MDD are above(below) the regression line and 23(41) HC are above(below) the regression line. The elevated Ridge Gene Age Gap Estimate (R-GAGE residual) in MDD compared to HC is statistically significant (Fig. 8, t-test P-value = 0.000186). Also note there is no association between MDD/HC and chronological age (t-test P-value=0.9756).

**Figure 7.**
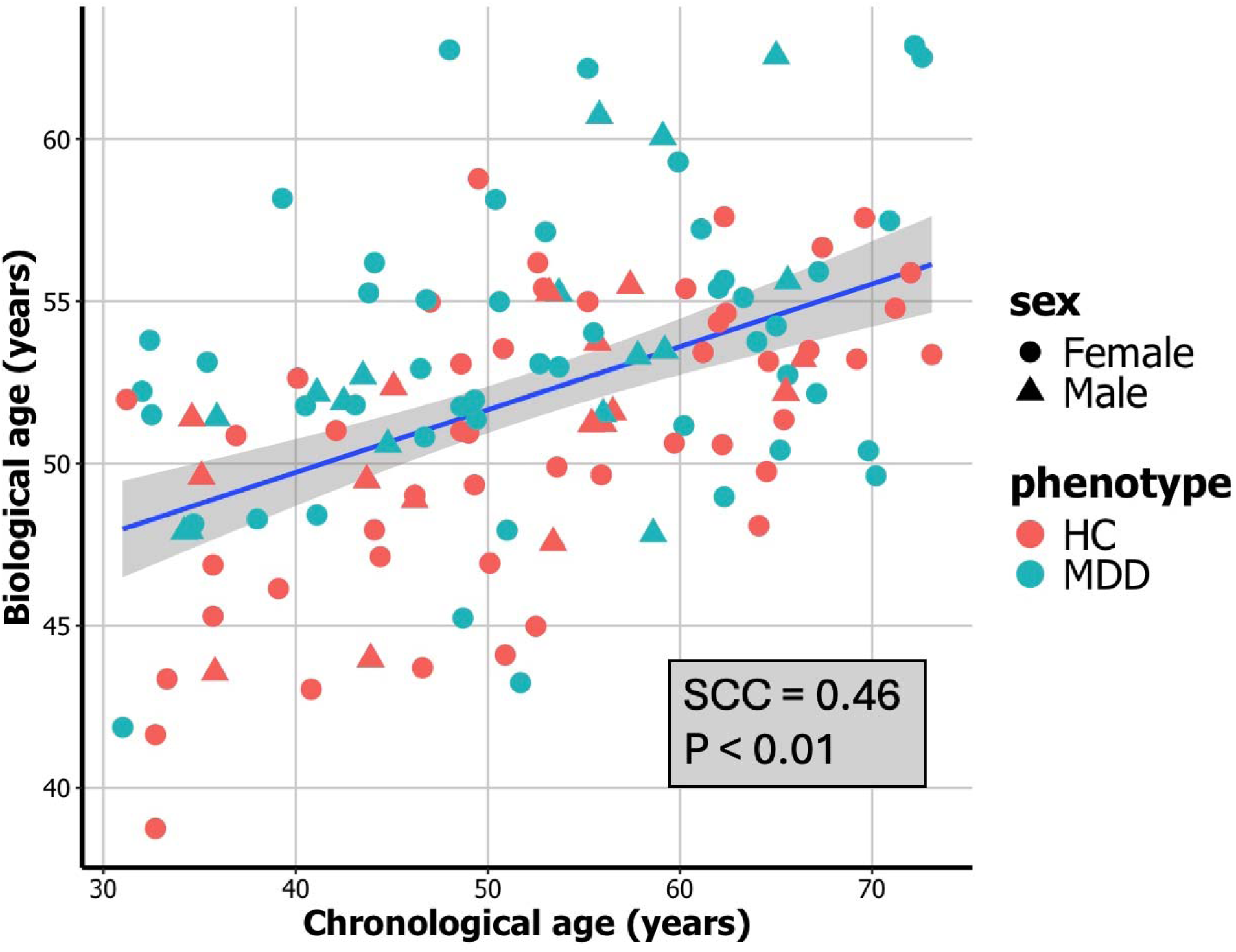
Gene age versus chronological age in replication microarray data (GSE98793). Gene age regression line (blue line, Spearman correlation coefficient (SCC) = 0.46, P-value < 0.01) is based on Ridge regression in the replication microarray data using only the 19 genes from the original L-GAGE model (21 genes less 2 missing). The points are colored by MDD (blue) and HC (red) and shaped by Female (circle) and Male (triangle). For MDD, 41(23) are above(below) the regression line and for HC, 23(41)are above(below) the regression line. The Ridge Gene Age Gap Estimate (R-GAGE) is higher for MDD than HC participants (see Fig. 8, t-test P-value = 0.0001856).

**Figure 8.**
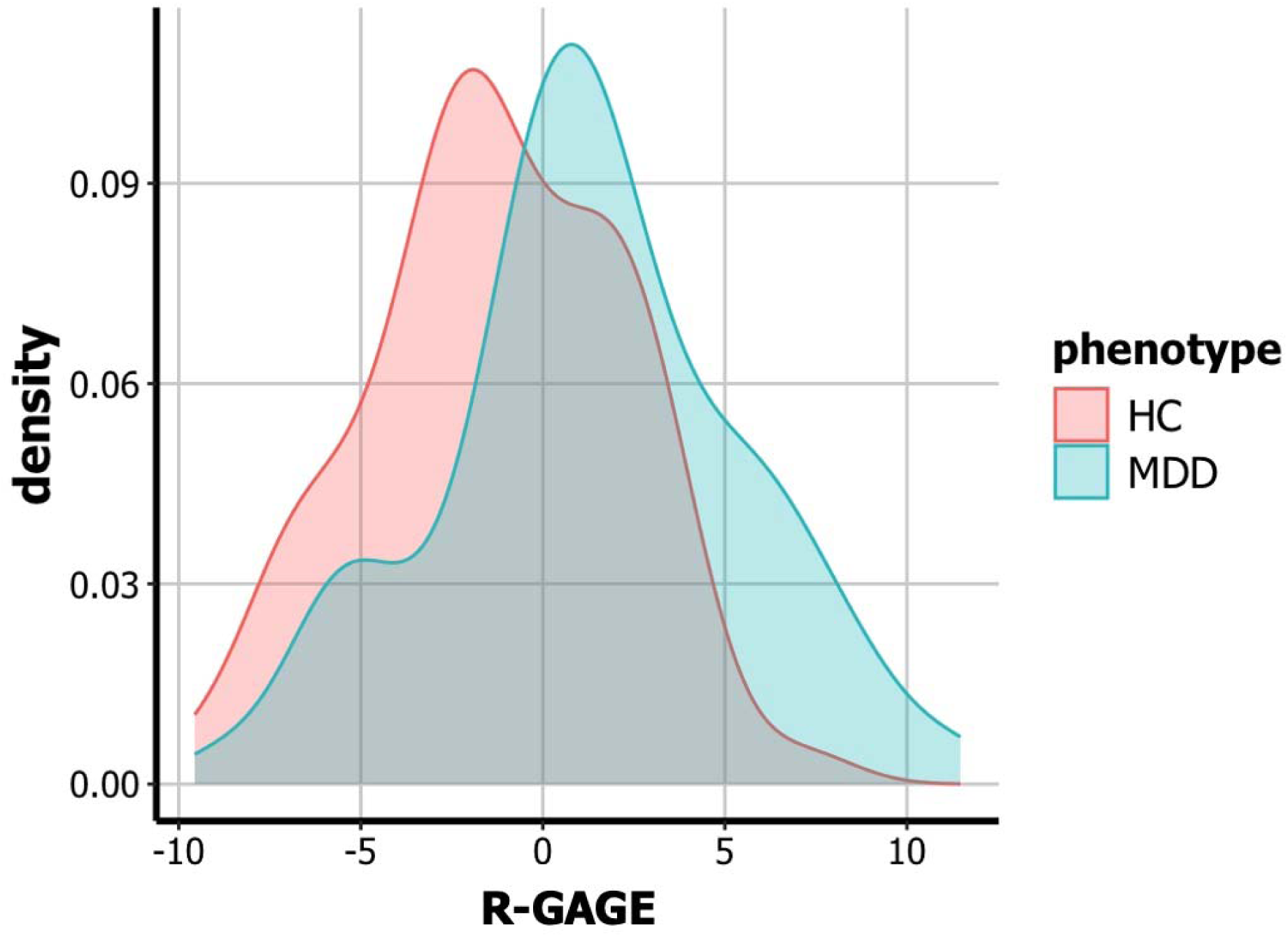
Density plot of the Ridge based Gene Age Gap Estimate (R-GAGE) in replication microarray data (GSE98793). The Ridge age model (Fig. 7) is trained on the replication microarray data using19 genes from the original L-GAGE model. A positive R-GAGE residual (x-axis) indicates a sample above the gene age regression line and negative below (Fig. 7). Biological age relative to chronological age (R-GAGE) is higher in MDD (blue) participants than in HC (red) (T-test P-value = 0.0001856).

## Conclusion

We presented a procedure for creating an expression-based biological age model using LASSO penalized regression, and we explored the association between the residual (the LASSO-based Gene Age Gap Estimate or L-GAGE) and MDD while adjusting for chronological age and sex. We found increased biological aging based on L-GAGE in MDD versus HC participants with an effect size similar to a previous study (Cole et al., 2021), but the difference was not statistically significant (see discussion below about replication). We found a statistically significant MDD-Age interaction for L-GAGE when age is dichotomized with threshold 40 years. We used multiple statistical criteria to verify this threshold based on the age distribution. Using a higher threshold results in a very sparse older group. The MDD-Age interaction could indicate an effect of lifetime number of MDD episodes on biological aging that is not detectable until middle-age. The interaction effect remained significant when adjusting for chronological age and sex, and we emphasize the importance of including age as a covariate in biological age association tests to avoid confounding due to regression to the mean (Le et al., 2018).

Elevated gene aging found previously in MDD participants was also observed in our data based on L-GAGE but was not statistically significant. However, we found evidence for the generalizability of the L-GAGE model by using the 21 LASSO-selected genes to test for elevated gene age in an independent microarray gene expression dataset (recall the discovery data used RNA-Seq). A limitation of this replication is the retraining of the model coefficients, which was necessary because only 19 of the 21 genes could be reliably mapped to microarray probes. Also, distribution differences between platforms changes the scale of the original regression coefficients. And so, while we did retrain the coefficients for microarray data, we restricted ourselves to the same genes selected in the discovery cohort, and we found a statistically significant elevation of gene age in MDD compared to HC. The age distributions are also somewhat different, with the discovery set skewing younger and the replication set skewing older. An interesting future work would be to integrate these data and other cohorts to get a broader age range with a gene age model with more generalizability.

We note a few of the top age-associated genes, such as CCNE1, NAA20, SESTD1, and TAF9, that have been associated with aging, senescence, and infectious disease. In a study of Lung Adenocarcinoma, CCNE1 gene expression was found to be correlated with patients’ age (Ullah et al., 2022), and NAA20 and SETD1A are involved in senescence, which is related to aging and age-related diseases. It was shown that depletion of NAA20 in non-transformed mammal cells led to senescence (Elurbide et al., 2023), and in another study knockdown of SETD1A triggered cellular senescence. (Tajima et al., 2019). TAF9 cross-reactivity was shown to be associated with immunity to CMV (human cytomegalovirus) in the context of autoimmune disease (Chen et al., 2021).

Pathway enrichment of a larger set of age genes, beyond the 21 L-GAGE genes, resulted in the detection of Infectious Disease, Adaptive Immunity, and SARS-CoV Infection pathways. As noted in (Cole et al., 2021), evaluating PBMC transcription can increase the risk for false positive immune pathways. Future work will involve pathway analysis based on other biological specimens and testing of the gene age model for MDD in RNA-seq data from postmortem brain areas. For example, the Stanley Medical Research Institute data includes frontal and cingulate cortex and hippocampus as well as several subcortical areas. Brain region-specific gene age effects on MDD could provide valuable insights into the etiology of MDD.

This study contributes a new approach to estimating biological aging and contributes to the evidence for the role of aging and inflammation in depression. Future studies are needed with broader age ranges, more uniform age distributions, large sample sizes, and utilization of MDD age-of-onset and number of depressive episodes. Future gene age models may help identify individuals that need different treatment or management for depression due to an increase in their relative biological age.

## Research data for this article

Data and code for this research are available at https://github.com/insilico/GeneAgeMDD.

